# Inertial Measurement Unit-Based Estimation of Foot Trajectory for Clinical Gait Analysis

**DOI:** 10.1101/595496

**Authors:** Yumi Ono, Koyu Hori, Hiroki Ora, Yuki Hirobe, Yufeng Mao, Hiroyuki Sawada, Akira Inaba, Satoshi Orimo, Yoshihiro Miyake

## Abstract

Gait analysis is used widely in clinical practice for the evaluation of abnormal gait caused by disease. Conventionally, medical professionals use motion capture systems or make visual observations to evaluate a patient’s gait. Recent biomedical engineering studies have proposed easy-to-use gait analysis methods involving wearable sensors with inertial measurement units (IMUs). IMUs placed on the shanks just above the ankles allow for the long-term monitoring of gait because the participant can walk with or without shoes during the analysis. As far as the authors know, there is no report of the gait analysis method that estimates stride length, gait speed, stride duration, stance duration, and swing duration at the same time. In this study, we tested a proposed gait analysis method that uses IMUs attached on the shanks to estimate foot trajectory and temporal gait parameters. We evaluated this proposed method by analyzing the gait of 10 able-bodied participants (mean age 23.1 years, nine men and one woman). Wearable sensors were attached to the participants’ shanks, and we measured three-axis acceleration and three-axis angular velocity with the sensors to estimate foot trajectory during walking. We compared gait parameters estimated from the foot trajectory obtained with the proposed method and those measured with a motion capture system. Mean accuracy (mean ± standard deviation) was –0.046 ± 0.026 m for stride length, –0.036 ± 0.026 m/s for gait speed, –0.002 ± 0.019 s for stride duration, –0.000 ± 0.016 s for stance duration, and –0.002 ± 0.022 s for swing duration. These results suggest that the proposed method is useful for evaluation of clinical gait parameters.

## 1 Introduction

Analysis of abnormal gait can provide important information about diseases. For example, patients with Parkinson’s disease (PD) often exhibit shuffling, festinating, and freezing of gait. The most widely used clinical rating scale for PD, the Unified Parkinson’s Disease Rating Scale, includes observation of gait (1). Patients with cerebellar disorders sometimes have a wide-based gait (atactic gait), and those with cerebral vascular disease sometimes exhibit a hemiplegia gait. Recent studies have also reported changes in gait, such as reduced gait velocity and stride length, in diseases with gait disorders and in other conditions such as Alzheimer’s disease (2) and depression (3).

Clinical gait analysis is performed mostly by health-care providers using visual observation (4). Although this method is the most readily accessible means of gait analysis available to health-care providers (5), it is a subjective and qualitative method that is inadequate for assessing changes in gait features during ongoing treatment interventions. It is also difficult for clinicians to share this information with health-care providers and patients. Motion capture systems are used widely in clinical gait analysis (6). Because they provide well-quantified and accurate results, these systems are at present considered to be the gold standard for clinical gait analysis (7). However, because the special equipment needed for motion capture is expensive and requires a large space, few medical institutions can use these systems for clinical gait analysis (5).

Several studies have proposed simple gait analysis methods using inertial measurement units (IMUs) to solve the problems described above (8–12). IMUs used in these methods are inexpensive and wearable. Sabatini et al. proposed an IMU-based gait analysis method that estimates a two-dimensional trajectory in the sagittal plane of a foot during walking (13). Some studies have proposed gait analysis methods that estimate the three-dimensional foot trajectory during walking in a stepwise manner to obtain values of foot clearance (14–16). The trajectory estimation methods reported in both Sabatini et al. (13) and Kitagawa and Ogihara (8) use an IMU attached on the dorsum of the foot and are better for obtaining this gait feature. Another proposed gait analysis method to estimate stride length uses an IMU attached on the shank for assessment of gait in people with PD but without estimating trajectory (17). However, this method can obtain stride length only and is not suitable for analyzing gait in people with diseases other than PD that cause gait abnormality, such as Alzheimer’s disease and depression.

In this study, we propose a novel gait analysis method for clinical purposes that uses IMUs attached on the shanks to estimate foot trajectory.

## 2 Proposed method

### 2.1 Sensors used and wearing method

Our proposed gait analysis system is illustrated in Figure 1A. For gait analysis, we used two IMUs (TSND121, ATR-Promotions, Kyoto, Japan; Figure 1B) with a triaxial accelerometer (±8 G range), triaxial gyroscope (±1000 degrees per range), and Android OS tablet (ZenPad10, ASUSTeK Computer Inc., Taipei, Taiwan; Figure 1C). Raw accelerometer and gyroscope signals are sampled at 100 Hz (16 bits per sample). The size of the IMU is 37 mm × 46 mm × 12 mm and its weight about 22 g. IMUs are attached on the shanks (just above the ankles) with bands (Figure 1B). The inertial coordinate system used to represent foot orientation and position relative to the global coordinate system is shown in Figure 1B. Acceleration and angular velocity data of both the shanks measured during walking are transmitted to the tablet through Bluetooth.

**FIGURE 1.**
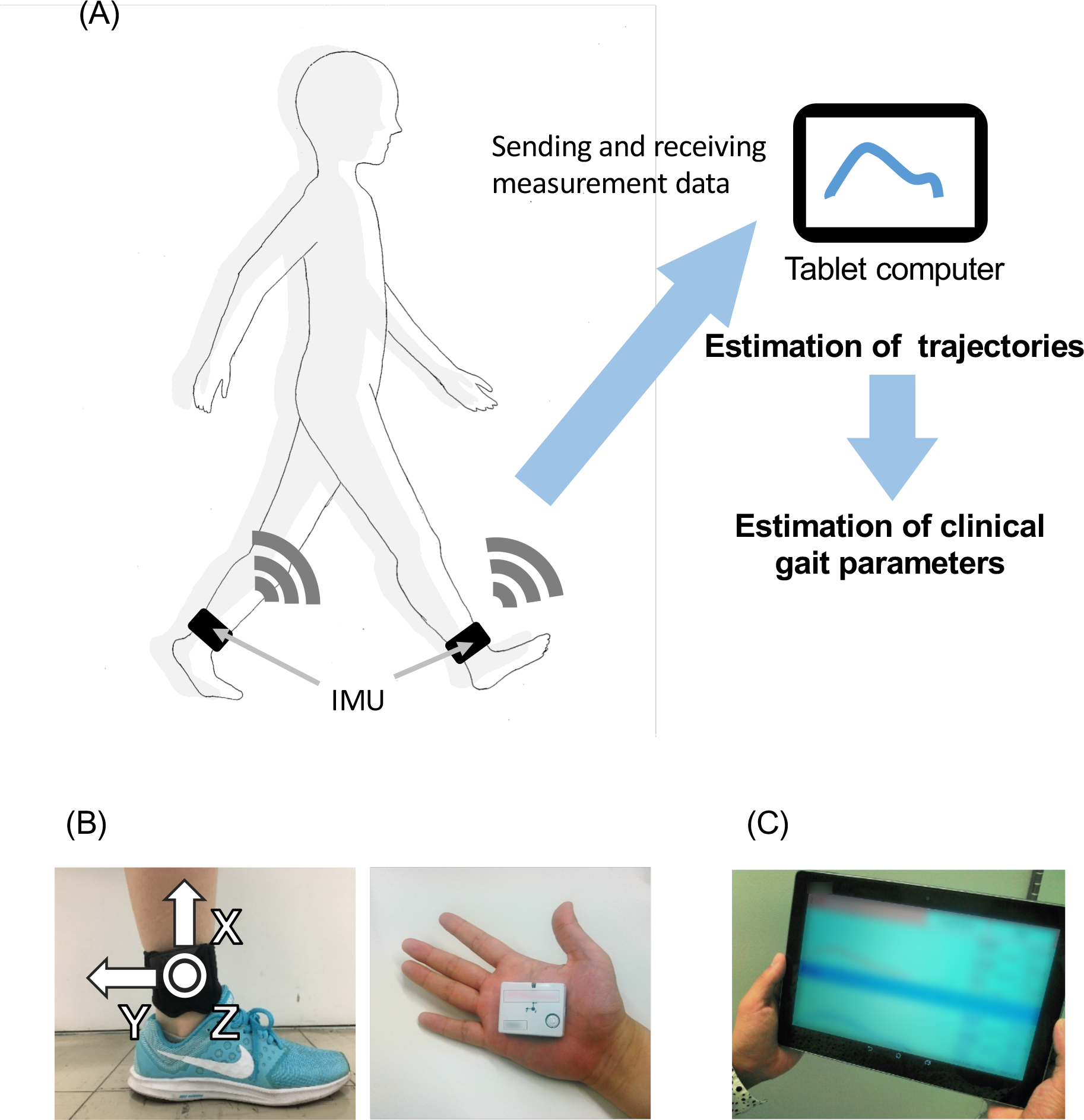
Overview of the system. **(A)** System configuration of the proposed method, including **(B)** a wearable sensor and its attachment on the shanks (just above the ankles), and **(C)** a tablet computer.

### 2.2 Algorithm for trajectory estimation

Our proposed method comprises two steps: dissociation of continuous gait data into multiple steps and three-dimensional trajectory estimation in a stepwise manner. Each process is described as follows.

### 2.3 Stepwise dissociation from angular velocity signals

We divide stepwise dissociation into four steps: (1) smoothing and finding the heel-strike and toe-off points; (2) finding the heel-strike and toe-off points; (3) quadratic regression; and (4) calculating the split point. One walking cycle is defined as a single step, and the starting point of each cycle is defined as the steadiest point between heel strike and toe-off. We identify the steadiest point in each cycle based on raw angular velocity data in the *z*-axis *ω_z_*.

#### 2.3.1 Smoothing and finding the local maximums

The raw data contain much noise, and a median filter (window length: 5) is first used to smooth the data. We find the local maximum *l_k_* that is larger than the threshold (200 degrees/s; Figure 2A).

**FIGURE 2.**
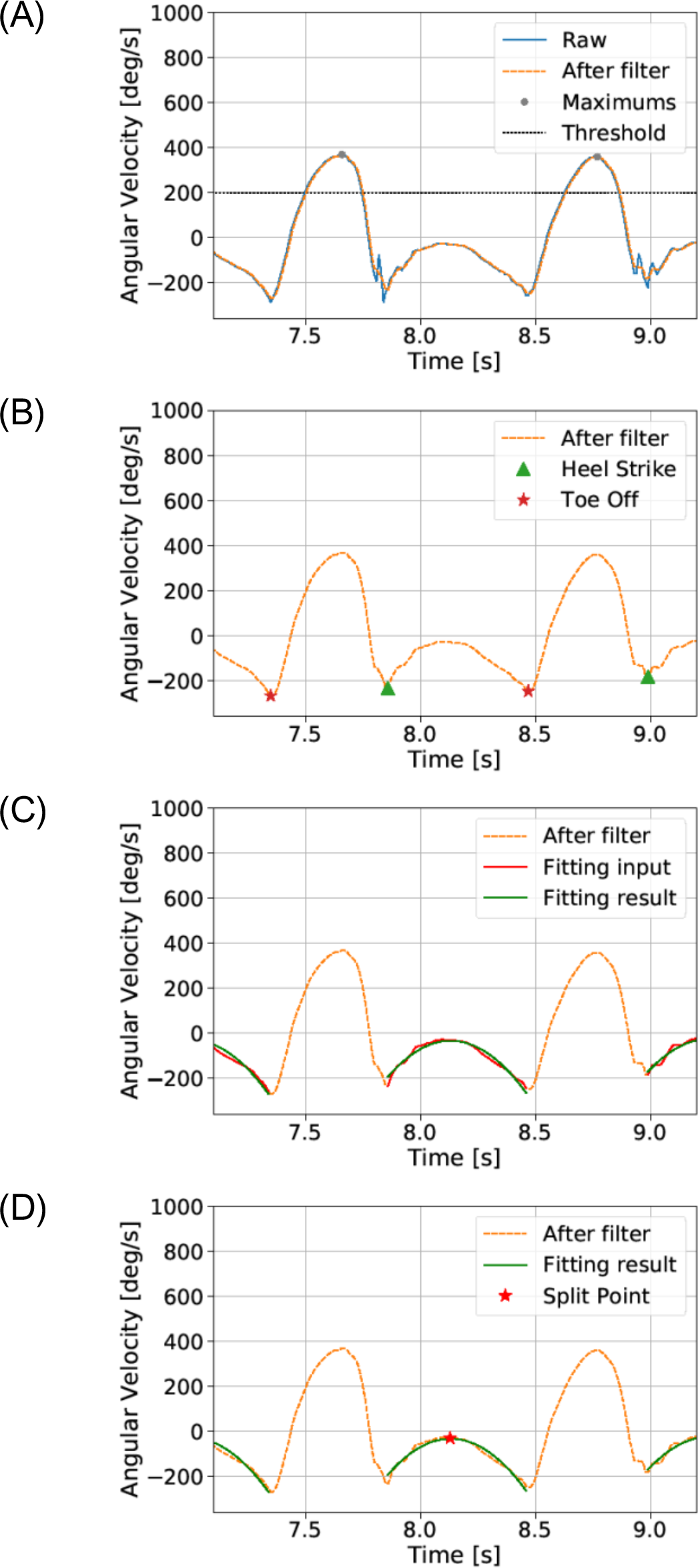
Dissociation of the continuous gait signal (see the section **Proposed method**). The raw data contain much noise, and a median filter is first used to smooth the data. **(A)** We find local maxima that are larger than a threshold and **(B)** then the heel-strike and toe-off points. **(C)** We assume a quadratic curve between the heel-strike point and the toe-off point. **(D)** Finally, the split point is defined as the maximum point of the quadratic fitting result. The gait cycle is defined by the data from the split point to the next split point.

#### 2.3.2 Finding the local minimums near the heel-strike and toe-off points

We then find the *k*-th local minimums near heel-strike *mhs_k_* and toe-off *mto_k_* points between the *l_k_* and *l_k_*_+1_. mhs_*k*_ is the local minimum closest to the right of *l_k_*, and *mto_k_* is the local minimum closest to the left of *l_k_*_+1_ (Figure 2B).

#### 2.3.3 Quadratic regression

We assume *ω_z_* between *mhs_k_* and *mto_k_* point to represent a quadratic curve (Figure 2C):

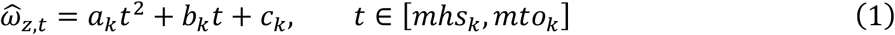

where 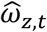 is the best-fitting result for angular velocity in the *z*-axis.

A quadratic regression is then used to fit the raw data and to calculate the parameters *a_k_*, *b_k_*, *c_k_*:

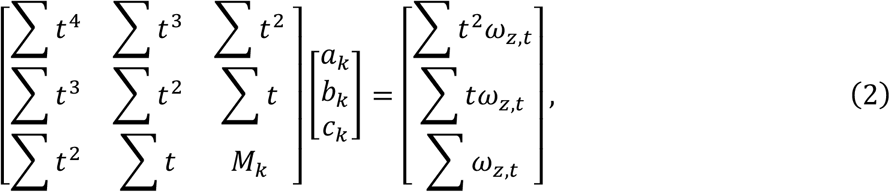

where *M_k_* is the number of points between *mhs_k_* and *mto_k_*.

#### 2.3.4 Calculating the split point

Finally, the segmentation point *sp_k_* is defined as the maximum point of the quadratic fitting result (Figure 2D):

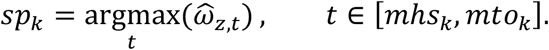

The *k*-th cycle is defined by the data between *sp_k_* and *sp_k+_*_1_. In each cycle, we estimate the trajectory.

### 2.4 Estimating the trajectory of a step

The foot trajectory in each cycle can be calculated by integrating the acceleration between each segmentation point. Two coordinate systems are applied: a laboratory coordinate system (*e*) and a sensor coordinate system (s). Because the raw data from the sensor are represented by the time-variant sensor coordinates, we need to transpose them into time-invariant laboratory coordinates. Acceleration in the laboratory coordinates can be converted from measured sensor acceleration using a rotational matrix:

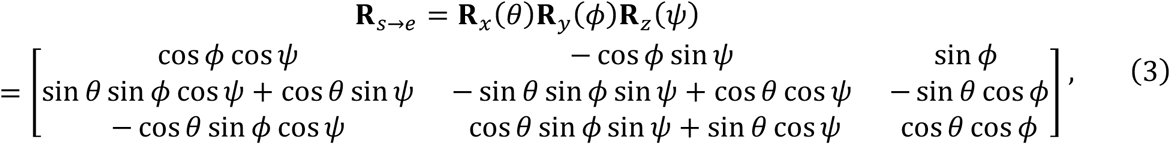

where *θ*, *ϕ*, and *ψ* are the Euler angles around the *x*-, *y*-, and *z*-axes.

We divide the estimation process of the foot trajectory into five steps: (1) calculate the initial Euler angles; (2) calculate the time derivative of the Euler angles; (3) calculate the Euler angles using the integral; (4) transpose the accelerations using a rotational matrix; and (5) calculate the trajectory using the double integral.

#### 2.4.1 Calculate the initial Euler angles

At the beginning of each cycle, we can assume that the foot is in full contact with the floor and is momentarily stationary. The accelerometer is assumed to detect only gravitational acceleration **g** at the outset:

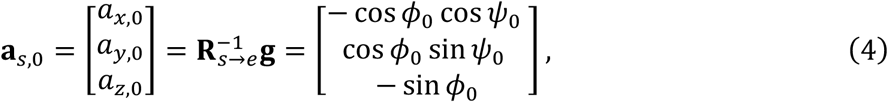

where **a**_*s*_,_0_ is the initial acceleration vector in the sensor coordinate, and *ϕ*_0_ and *ψ*_0_ are the initial Euler angles around the *y*- and *z*-axes.

Therefore, the initial Euler angles vector θ_0_ can be calculated as:

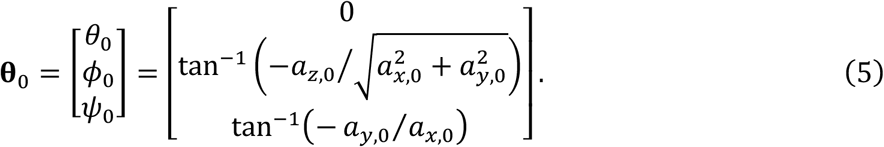

#### 2.4.2 Calculate the time derivative of the Euler angles

The relation between the *i*th angular velocity ω_*i*_ from the gyroscope and the time derivation of Euler angles 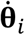 can be calculated as:

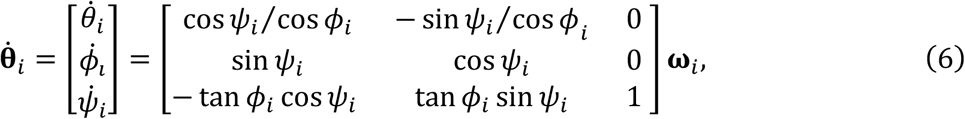

where *ϕ_i_* and *ψ_i_* are the *i*th Euler angles around the *y*- and *z*-axes.

#### 2.4.3 Calculate the Euler angles by integral

Euler angles are derived by the integration of Euler angle derivation:

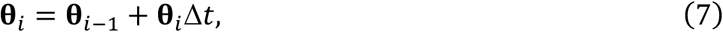

where Δ*t* is the sampling rate.

#### 2.4.4 Transpose accelerations using a rotational matrix

The rotational matrix calculated by equation (3) is used to estimate the acceleration in laboratory coordinates **R**_*S*→e_**a**_*S*_, which contains the gravitational acceleration. Linear acceleration **a**_*e*_ in laboratory coordinates can then be calculated by simply subtracting gravity:

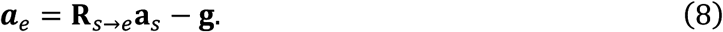

#### 2.4.5 Calculate trajectory using the double integral

The *i*-th velocity **v**_*e,i*_ in the laboratory coordinates is estimated by integration of linear acceleration **a**_*e,i*_:

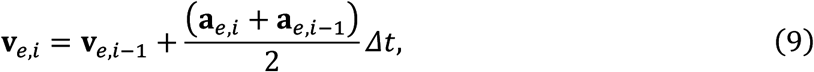

and the *i*-th foot trajectory **p**_*e,i*_ is estimated by integration of v_*e,i*_:

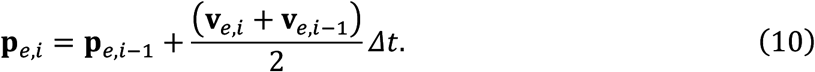

### 2.5 Reduction of Brownian noise

Integration would drift because of the IMU sensors error, and the calculations of velocity and trajectory are corrected by the constraint condition. In each cycle, both the initial value and the end value of the velocity in three directions and the trajectory in the vertical direction can be assumed as 0. The algorithm below shows how to estimate velocity.

First, the forward integral and back integral are calculated separately from linear acceleration **a**_*e,i*_:

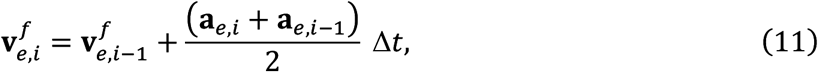

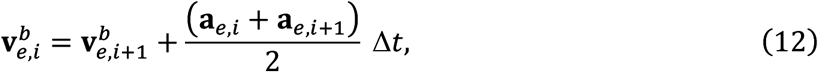

where the superscripts *f* and *b* mean forward and backward, respectively. The correction result can then be derived by the weighted average of the forward and backward integral:

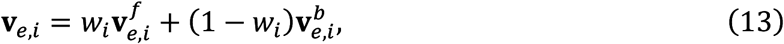

where *w_i_* is the weight and *w_i_* ∈ [0,1].

Because 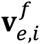 is more accurate near the starting point and 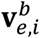 is more accurate near the end point, the function for calculating *w_i_* should increase monotonically. Here, we choose the sigmoid function to calculate *w_i_*:

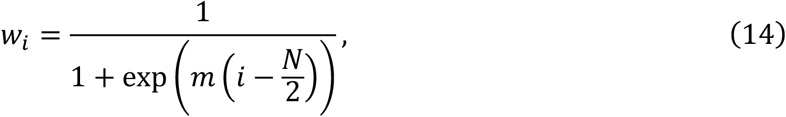

where *N* is the number of points in the current cycle and *m* is a hyperparameter calculated from experiment, which we choose as *m* = 0.1. The trajectory in the vertical direction can also be calculated using the above algorithm.

### 2.6 Estimation of spatial and temporal parameters for clinical gait analysis

The gait events included the heel-strike (HS) and toe-off (TO) were extracted first. The HS and TO events detection algorithm was based on the peak detection of the raw angular velocity in sagittal plane ω_*z*_. At the end of the swing period, several of negative peaks can be observed in ω_*z*_ and the first one is associated with the HS instant (18). Prior of the swing period, a negative peak is associated with the TO instant (18).

For the definition of each gait event search region, a method that utilized the variation pattern of shank tilt angle which inspired from a instep-based previous study (19) was introduced. At the end of the swing period, the shank will rotate counterclockwise around the knee and reach the maximum forward. Then, the clockwise rotation start and the foot contact the ground to produce the HS instant. In this process, *θ_z_* will appear to increase first and then decrease thus a positive peak will appear in *θ_z_* before the HS instant where *θ_z_* was computed via the integration of *ω_z_* with sampling interval Δt. For the convenience of description, we refer to this instant where peak occurrence as shank-max-forward (SMF). Similarly, after the TO instant, the ankle will slightly lift and rotate counterclockwise until reaching a certain height. Then it will start to rotate counterclockwise and present a negative peak in *θ_z_*. We refer this instant as shank-max-backward (SMB) for convenience. As a result, SMF and SMB of *θ_z_* can be used to define a proper search interval of the HS and TO. We found the SMF and SMB via a peak search algorithm *signal.find_peaks* in SciPy (v1.2.0) which can find proper peaks via the prominence (define intrinsic height of a peak) and the distance (define the distance between peaks) properties. Then, SMFs are the positive peaks and SMBs are the negative peaks whose prominence is larger than 0.2 [rad] and the distance is at least 0.4 [s]. Finally, HS is defined as the first peak appears after each SMF instant, and the TO is the minimum of angular velocity in the interval (SMB — 0.3 [s], SMB).

Estimated stride length *SL* is calculated by the trajectory in the *y* and *z* direction:

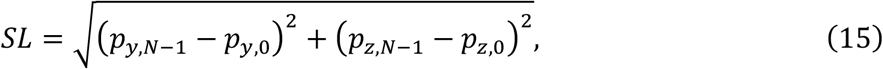

where **p**_*e,i*_ = (*P*_*x,i*_,*P*_*y,i*_,*P*_*z,i*_)^*T*^. Estimated stride duration is defined as the time from one heel strike to the next heel strike. Estimated gait speed is defined as the value obtained by dividing stride length by stride duration. Estimated stance duration is defined as the time from heel strike to toe-off of just before the next heel strike. Estimated duration time is defined as the time from toe-off to the next heel strike.

## 3 Evaluation of the proposed method

### 3.1 Overview of the experimental evaluation

We conducted an experimental evaluation of our proposed method to validate the accuracy of the trajectory estimation of the shanks (just above the ankles) to verify whether it allows the analysis of gait for clinical purposes. Ten healthy people participated in the experiment, and we used motion capture systems as the gold standard for the assessment of gait in the clinical setting. We evaluated the accuracy of our proposed method for calculating the estimated trajectory and clinical gait parameters. We used an IMU attached on the shanks for the experimental evaluation.

An optical motion capture system (Nobby Tech. Ltd., Tokyo, Japan) was used as the reference system. We used 12 cameras, and the motion capture volume was about 2 m × 7 m × 1 m (Figure 3A). The position error of the markers of the motion capture system was less than 1 mm. Three markers were attached on each foot as shown in Figure 3B. Two of three markers were attached on the heel and the toe for assessment of gait parameters. The third marker was attached on the IMU for evaluation of the trajectory estimation.

**FIGURE 3.**
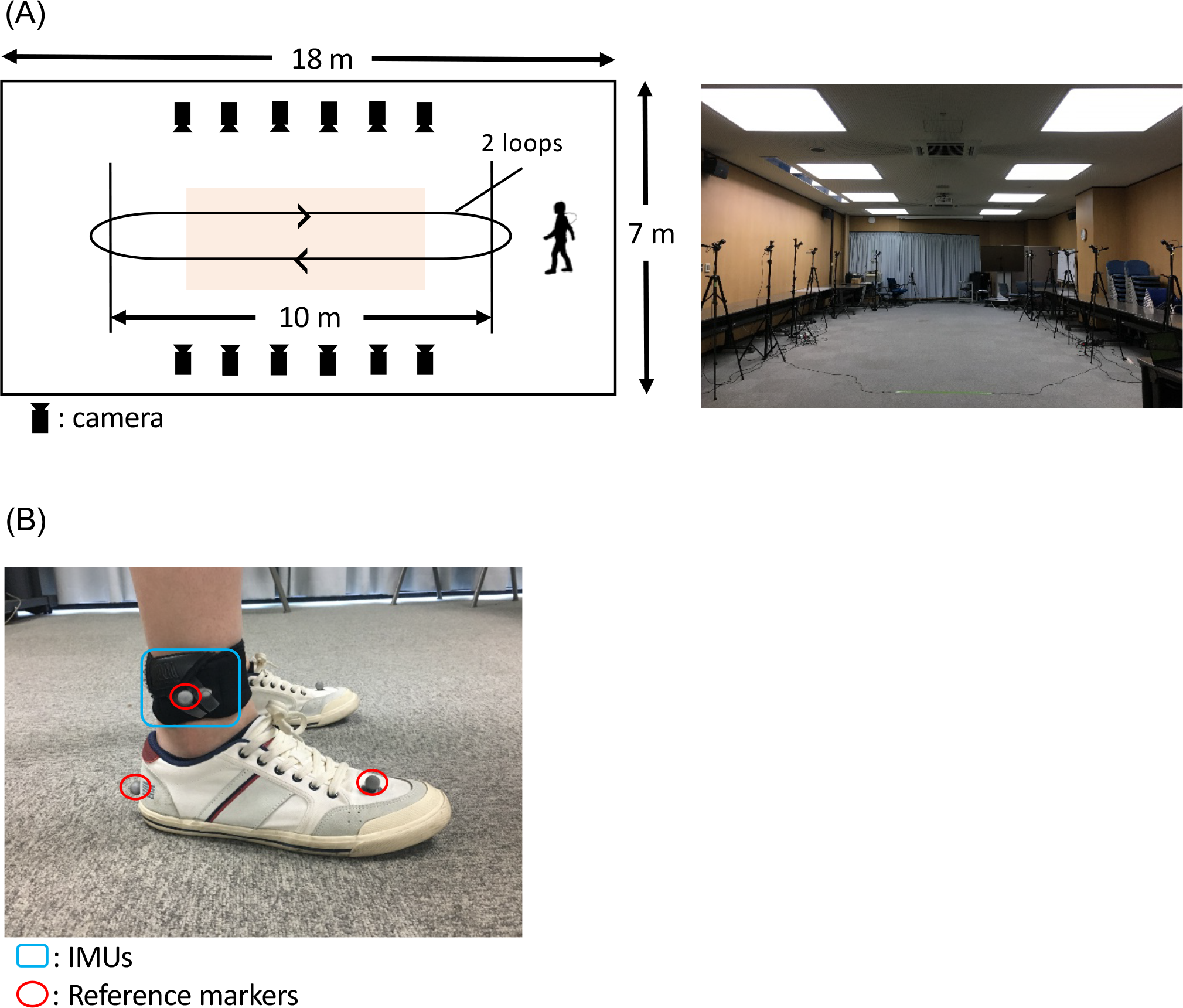
Protocol for the experimental evaluation of the proposed method. **(A)** A room at Tokyo Institute of Technology was used (**A**, right). The orange-shaded area shows the measurement area for gait analysis. The participants with IMUs and optical markers **(B)** walked around the room twice, as indicated by the arrows (**A**, left).

### 3.2 Participants

Ten healthy participants (mean age 23.1 years, nine men and one woman), with no history of gait abnormalities, were recruited from Tokyo Institute of Technology. The characteristics of each subject are summarized in Table 1. The Ethics Committee of Tokyo Institute of Technology approved the protocols for this study, and all participants provided written informed consent.

**TABLE 1.**
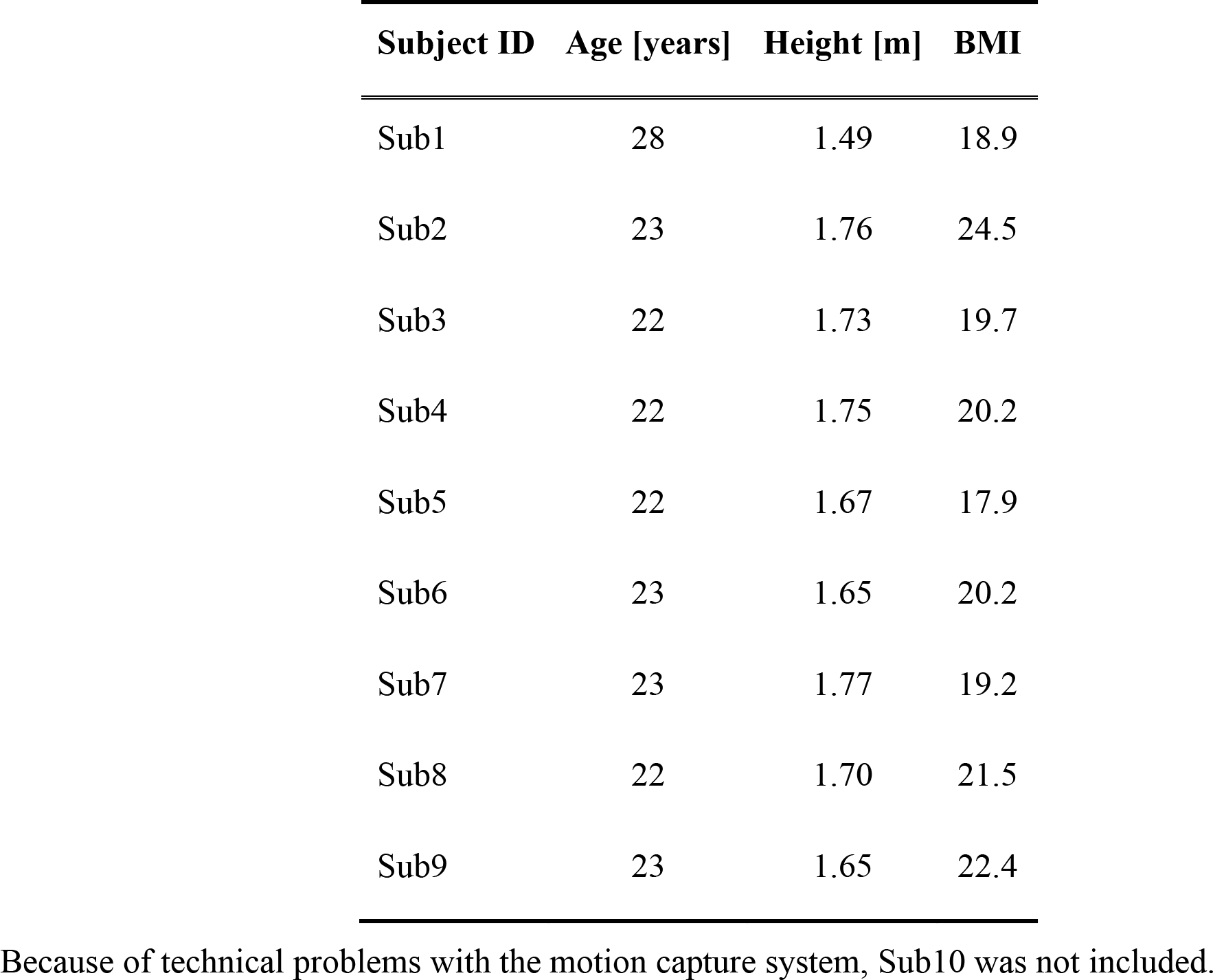
Summary of participant characteristics.

| **Subject ID** | **Age [years]** | **Height [m]** | **BMI** |
| --- | --- | --- | --- |
| Sub1 | 28 | 1.49 | 18.9 |
| Sub2 | 23 | 1.76 | 24.5 |
| Sub3 | 22 | 1.73 | 19.7 |
| Sub4 | 22 | 1.75 | 20.2 |
| Sub5 | 22 | 1.67 | 17.9 |
| Sub6 | 23 | 1.65 | 20.2 |
| Sub7 | 23 | 1.77 | 19.2 |
| Sub8 | 22 | 1.70 | 21.5 |
| Sub9 | 23 | 1.65 | 22.4 |

Because of technical problems with the motion capture system, Sub10 was not included.

### 3.3 Experimental task

Each participant walked two trials on a flat floor at his or her own self-selected natural pace and a slow pace. Each participant walked back and forth across the room twice during each trial as instructed. We used the gait data obtained as the participants walked in the central area of the room.

### 3.4 Validation of the location of IMUs

To consider the validity of the estimation of foot trajectory from IMUs attached on the shanks, we calculated correlations between stride length estimated with the proposed method, measured with a motion capture marker attached on the IMU, and measured with a motion capture marker attached on the heel.

### 3.5 Application of the proposed method to patients with gait disorders

We used the proposed method to analyze gait in one healthy elderly participant and four PD patients. The healthy elderly participant was recruited from a public interest incorporated association that provides human resource services for elderly people and is located in Machida City, Tokyo. PD patients were recruited from Kanto Central Hospital, Tokyo. PD patients had been diagnosed by a doctor. The exclusion criteria for this study were past history of other neurological or orthopedic disorders that can affect gait or posture (excluding PD). The participants provided written informed consent in accordance with the Ethics Committee of Tokyo Institute of Technology. The Kanto Central Hospital Ethics Committee and The Ethics Committee of Tokyo Institute of Technology approved the protocol for this study.

## 4 Results

To evaluate the proposed method, the shank (just above the ankle) trajectory and clinical gait parameters calculated by our proposed method were compared with those collected by the motion capture system. The comparisons of trajectory information were conducted in the sagittal plane. Six clinical gait parameters were compared. Because of technical problems with the motion capture system, the data for one of the 10 participants were excluded from the analyses.

### 4.1 Comparison of trajectory estimated with the proposed method and the motion capture system

The trajectories of our proposed method and the reference data are shown in Figure 4. The R value between displacement in the direction of forward movement calculated with the proposed method and measured with a marker attached on the IMU was 0.991 (Figure 5A). The R value between the maximum vertical displacement calculated with the proposed method and measured with a marker attached on the IMU was 0.926 (Figure 5A).

**FIGURE 4.**
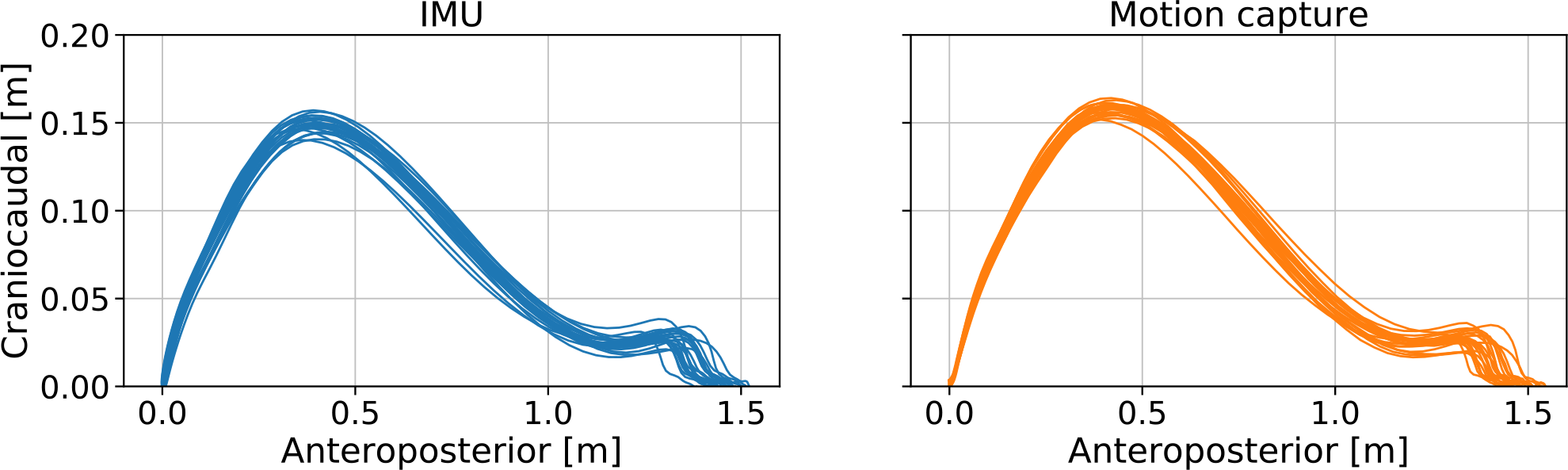
Errors between the estimated foot trajectory identified with our proposed method and reference data from the motion capture system projected in the **(A)** sagittal and **(B)** horizontal planes.

**FIGURE 5.**
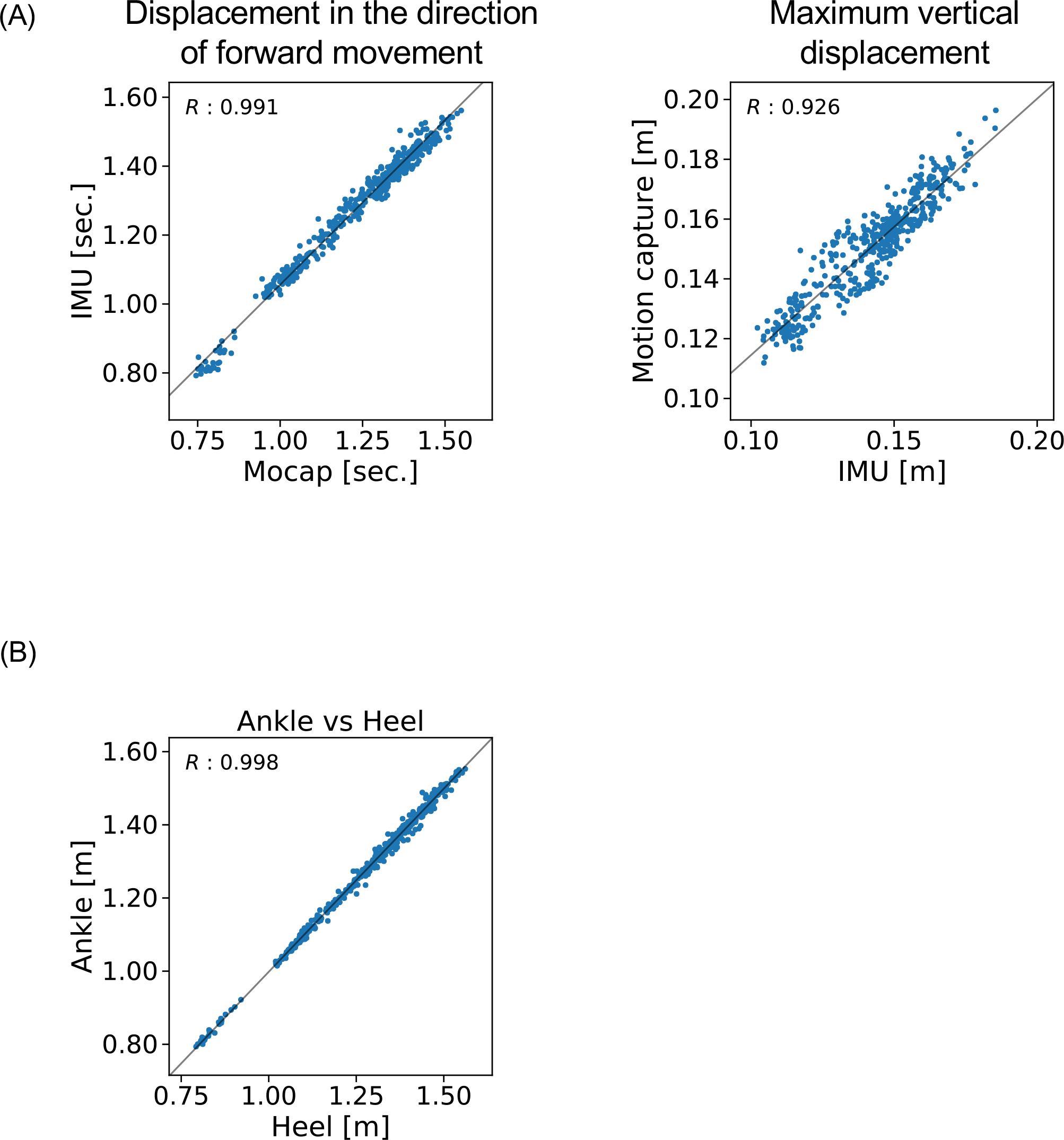
R values for. **(A)** the displacement in the direction of forward movement and maximum vertical displacement estimated with the proposed method and **(B)** measured with a marker attached on the IMU and measured with a marker attached on the heel.

### 4.2 Validation of the location of IMUs

The R value between displacement in the direction of forward movement calculated with the marker attached on the IMU and that measured with the marker attached on the heel was 0.998 (Figure 5B).

### 4.3 Estimation of clinical gait parameters

The means and standard deviations of the gait parameters compared between the proposed method and the motion capture system are summarized in Table 2. The mean error of stride length was −0.046 ± 0.026 m (Figure 6A). The R value between displacement in the direction of forward movement calculated with the proposed method and measured with a marker attached on the heel was 0.991 (Figure 6A). The mean error values were as follows: −0.036 ± 0.026 m/s for gait speed (Figure 6B); −0.002 ± 0.019 s for stride duration (Figure 6C); −0.000 ± 0.016 s for stance duration (Figure 6D); and −0.002 ± 0.022 s for swing duration (Figure 6E).

**TABLE 2.** Comparison between the results of the proposed method and a motion capture system.

| **Parameter** | **Difference** |  |
| --- | --- | --- |
|  | **Mean** | **SD** |
| Stride length [m] | −0.046 | 0.026 |
| Gait speed [m/s] | −0.036 | 0.029 |
| Stride duration [s] | −0.002 | 0.019 |
| Stance duration [s] | 0.000 | 0.016 |
| Swing duration [s] | −0.002 | 0.022 |

**FIGURE 6.**
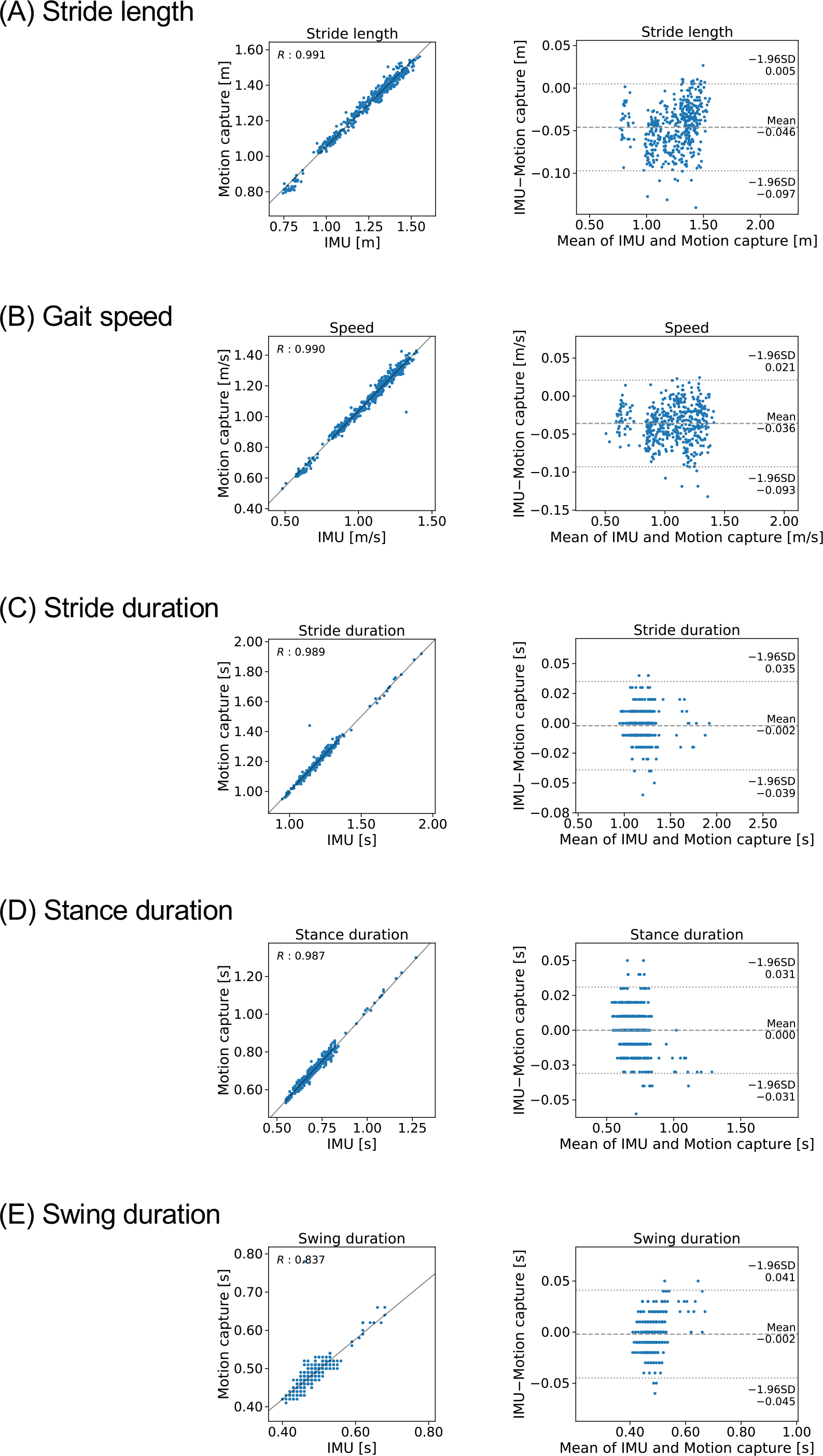
Comparison of the proposed method and the gold standard in stepwise manner. Mocap, motion capture. Scatter plots and, Bland and Altman plots of **(A)** stride length, **(B)** gait speed, **(C)** stride duration, **(D)** stance duration, and **(E)** swing duration.

### 4.4 Application of the proposed method to patients with a gait disorder

The shank trajectory over 15 steps for each participant is shown in Figure 7. The mean clinical gait parameters of the PD patients are summarized in Table 3.

**TABLE 3.** Mean clinical gait parameters of PD patients.

| **Patient ID (mH&Y)** |  | **Pt1 (2)** | **Pt2 (2)** | **Pt3 (4)** | **Pt4 (4)** |
| --- | --- | --- | --- | --- | --- |
|  | Stride length [m] | 1.03 (0.04) | 1.14 (0.04) | 0.29 (0.03) | 0.52 (0.05) |
|  | Gait speed [m/s] | 0.84 (0.04) | 1.05 (0.04) | 0.30 (0.02) | 0.51 (0.06) |
| **Parameter** | Stride duration [s] | 1.22 (0.04) | 1.09 (0.02) | 0.97 (0.06) | 1.02 (0.05) |
|  | Stance duration [s] | 0.71 (0.04) | 0.57 (0.03) | 0.62 (0.05) | 0.62 (0.06) |
|  | Swing duration [s] | 0.52 (0.04) | 0.52 (0.02) | 0.35 (0.04) | 0.40 (0.04) |

Values are expressed as mean (standard deviation). mH&Y=modified Hoehn and Yahr scale.

**FIGURE 7.**
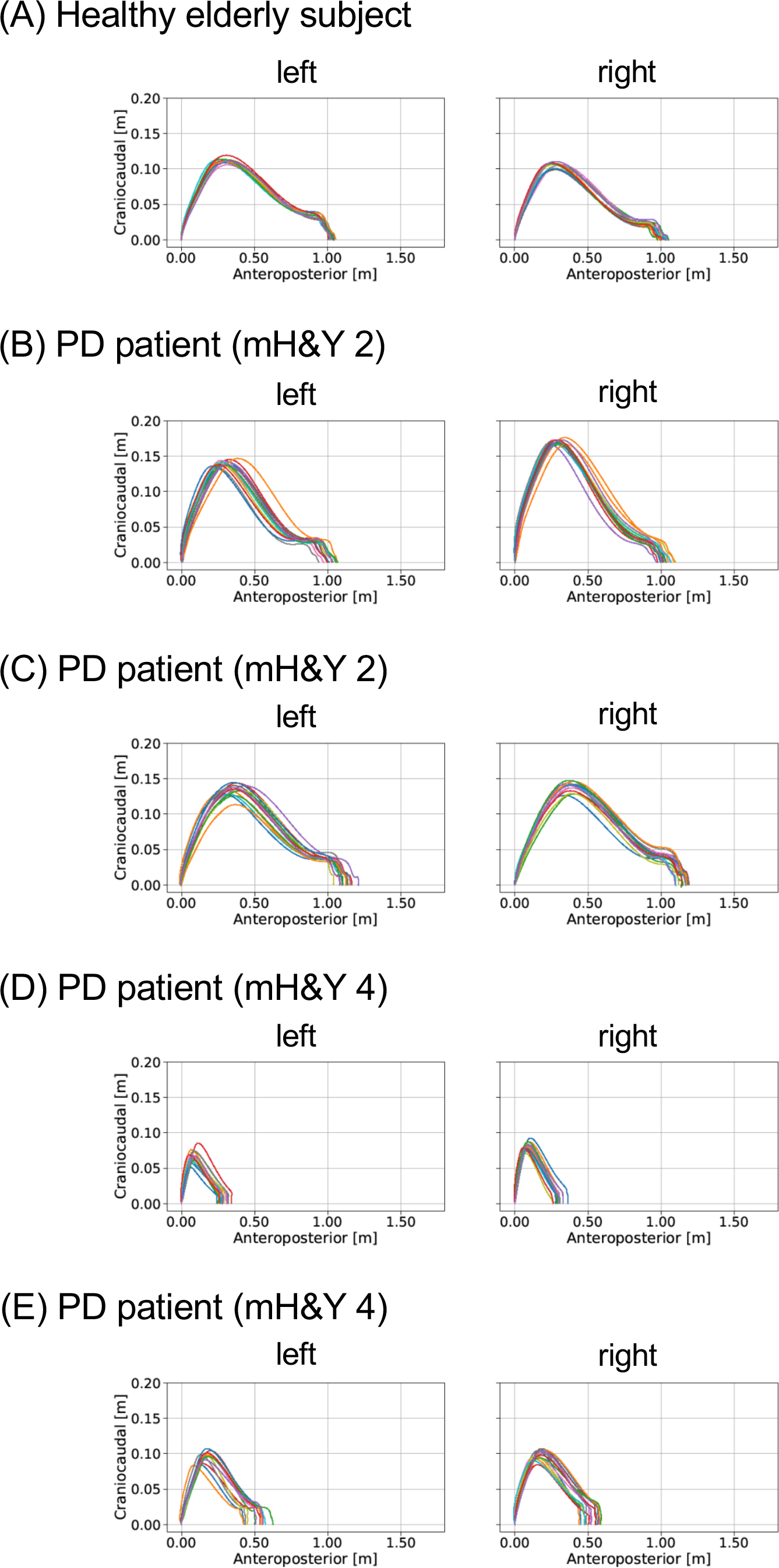
Examples of the application of our proposed method for analyzing gait in patients with PD. mH&Y, modified Hoen and Yahr scale. The estimated trajectories of **(A)** healthy elderly subject, **(B)** PD patient (mH&Y 2), **(C)** PD patient (mH&Y 2), **(D)** PD patient (mH&Y 4), and **(E)** PD patient (mH&Y 4) were plotted.

## 5 Discussion

We have proposed a new method for gait analysis that uses IMUs attached on the shanks to estimate foot trajectory. The experimental results show that the proposed method could be used to calculate clinical gait parameters by estimating foot trajectory.

### 5.1 The proposed system and method

The proposed gait analysis method comprises two IMUs with a triaxial accelerometer, triaxial gyroscope, and tablet computer. This method can be applied in a variety of locations and is less expensive than conventional gait analysis methods such as motion capture systems. The clinical advantage is that the patient burden is low because of the light weight (about 24 g) and easy attachment of the IMUs. We therefore anticipate that the proposed method would be suitable for clinical gait analysis.

The R values between displacement in the direction of forward movement measured with the marker attached on the IMU and measured with the marker attached on the heel (0.998) indicate that the location of the IMUs is valid for estimating the stride length.

### 5.2 Accuracy of estimation of foot trajectory

The R value between displacement in the direction of forward movement estimated by the proposed method and measured with the marker of the motion capture system attached on the IMU indicates that displacement in the direction of forward movement estimated by the proposed method explained 98% of the variation in displacement in the direction of forward movement measured with the motion capture system.

### 5.3 Accuracy of estimation of clinical parameters

The mean error of stride length estimated with the proposed method was −0.046 ± 0.026 (Table 2). This result suggests that the proposed method can estimate clinical gait parameters such as stride length and may be applicable in clinical practice. Many studies (20, 21) have reported that stride length is shorter in PD patients than in healthy controls. For example, Morris et al. reported that stride length in PD patients in the off state was 0.96 ± 0.19 m, which was shorter than the stride length of 1.46 ± 0.08 m measured in healthy age-matched controls (22). Our findings suggest that the proposed method has sufficient accuracy for evaluating stride length in people with PD.

The R value between displacement in the direction of forward movement estimated by the proposed method and measured with the marker of the motion capture system attached on the heel indicates that stride length estimated by the proposed method explained 98% of the variation measured with the motion capture system. The result suggests that IMUs are potentially useful in clinical gait analysis.

### 5.4 Future work

We expect that further development of this method or other methods will enable us to evaluate quantitatively the effects of drugs and interventions such as rehabilitation in patients with gait disorders. In the future, we plan to assess patients with gait abnormalities, such as that caused by PD. We will validate the proposed method to determine whether it can identify abnormal gait patterns, including shuffle, short-steppage, and hemiplegia gaits.

## 6 Conclusion

Our results suggest that the proposed method is suitable for clinical gait analysis. This method can be used in a variety of locations, such as in the corridor of a medical center, unlike methods that use motion capture systems. Our proposed method is expected to enable clinicians to share objective information about gait features with health-care providers and patients.

## 8 Author Contributions

KH, YO, YH, YMa, HO, SO, and YMi designed the research; YO, KH, YH, HS, and AI performed the experiment; YO, HO, KH, YMa, and YMi analyzed the data; HO, YO, and YMi wrote the paper.

## 10 Acknowledgments

We thank Mr. Masatoshi Seki for assistance.

